# Experimental increase of worker diversity benefits brood production in ants

**DOI:** 10.1101/2021.01.05.425457

**Authors:** Marina N. Psalti, Dustin Gohlke, Romain Libbrecht

## Abstract

The reproductive division of labor of eusocial insects, whereby one or several queens monopolize reproduction, evolved in a context of high genetic relatedness. However, many extant eusocial species have developed strategies that decrease genetic relatedness in their colonies, suggesting some benefits of the increased diversity. Multiple studies support this hypothesis by showing positive correlations between genetic diversity and colony fitness, as well as finding effects of experimental manipulations of diversity on colony performance. However, alternative explanations could account for most of these reports, and the benefits of diversity on fitness in eusocial insects still await validation. In this study, we experimentally increased worker diversity in the ant *Lasius niger* while controlling for typical confounding factors. We found that experimental colonies composed of workers coming from three different source colonies produced more larvae and showed more variation in size compared to groups of workers coming from a single colony. We propose that the benefits of increased diversity stemmed from an improved division of labor. Our study confirms that worker diversity enhances colony performance, thus providing a possible explanation for the evolution of multiply mated queens and multiple-queen colonies in many species of eusocial insects.

## Introduction

Genetic relatedness plays an important role in the evolution of altruistic behaviors in animals^1^. Extreme altruism is found in colonies of eusocial Hymenoptera (ants, bees and wasps), where the workers forgo their own reproduction to help the queens produce offspring^2^. Such reproductive division of labor evolved in a context of high genetic relatedness, with a single female reproductive mated with a single male^3,4^. Most extant eusocial Hymenoptera species are still characterized by high genetic relatedness^3^.

Other species evolved colonies with lower relatedness among individuals, and thus higher genetic diversity^5^. In these species, colonies have one multiply mated queen and/or multiple queens. Prominent examples include the honeybee *Apis mellifera*, where queens can mate with up to 20 males^6–9^, and the Argentine ant *Linepithema humile*, where colonies may contain up to 60 queens^10^. There are costs associated with such strategies. Multiple mating increases risk of disease or predation, and requires more energy to locate the sexual partners and copulate^11,12^. Having multiple queens per nest lowers relatedness in the worker force and may favor the emergence of conflicts among workers, which could affect colony performance^13–17^.

The evolution of such strategies to increase genetic diversity in some eusocial insect species shows that they must have benefits in certain ecological conditions^18^. The potential benefits of increased genetic diversity include increased resistance to parasites via improved social immunity^19–25^ and a more efficient behavioral division of labor among workers^26–31^.

Behavioral division of labor is the repartition of tasks in the worker force. For example, some workers tend to stay inside the nest to nurse the brood while others forage for food. These tendencies likely stem from workers differing in their internal response threshold to perform specific tasks^32^. This response threshold is determined by a combination of extrinsic and intrinsic factors, such as the social environment, location in the nest, morphology, age, individual experience and genetic background^33–39^. The increasing evidence for genetic effects on worker behavior and division of labor^27–31,40–42^ is consistent with the hypothesis that increased genetic diversity in the worker force would result in a larger variation in threshold responses, and thus a more efficient division of labor, which in turn could increase fitness.

Several lines of evidence suggest that intracolonial genetic diversity increases fitness. The reports of such findings fall in one of three categories. First, there are theoretical studies that supported a link between diversity and performance^17,43–45^. Second, there are reports of correlations between genetic diversity and one or several fitness correlates in several species of bees, wasps and ants^26,46–54^. Third, there are reports of experimental manipulations of genetic diversity that affected colony performance, mostly in bees^19,20,23,24,36,48,55–58^.

However, there is still debate over whether increased genetic diversity directly enhances colony fitness^59^. First, finding correlations between genetic diversity and fitness components does not imply causation, and other correlative studies did not detect such an association^46,50,60–62^. Then, the strategy of many studies that experimentally manipulated genetic diversity was to decrease it in species with naturally high diversity. For example, in the highly polyandrous honey bee, the artificial insemination of queens with the sperm from a single male reduced the performance of their colonies compared to queens inseminated with the sperm from multiple males^36,48,58^. In these studies, the decrease in colony performance associated with the low diversity treatment could be explained by the stress associated with not being in the natural state. Two studies in *Bombus terrestris* showed some benefits of artificially increased genetic diversity in a species with naturally lower diversity, but mostly in terms of resistance to pathogens^56,57^. Finally, experiments based on artificial insemination cannot disentangle between direct effects of genetic diversity among workers produced by the artificially inseminated queen and indirect maternal effects via the queen (e.g., on the number and quality of eggs produced) in response to the insemination with variable sperm diversity.

One way to get around the confounding maternal effects is to directly manipulate the diversity in the worker force. This experimental approach, so far restricted to the study of the effect of worker diversity on social immunity, showed an increased resistance to parasites and pathogens in bumble bees^20^ and ants^23^. Finally, experimental manipulations of worker size diversity in the ant *T. nylanderi* did not influence colony performance and fitness^63,64^.

To validate the effect of worker diversity on colony fitness in eusocial insects, there is a need for studies to go beyond the correlative approach, to experimentally increase worker diversity in species with naturally lower diversity, while controlling for potential maternal effects. Here, we use the black garden ant *Lasius niger* to experimentally produce colonies composed of workers from either one (low diversity) or three (high diversity) source colonies. These experimental colonies were then provided with a single, unrelated queen, and brood production was monitored over time. We report an increased brood production in experimental colonies with a more diverse worker force, thus showing that worker diversity enhances colony performance.

## Methods

### *Lasius niger* as a study system

To manipulate genetic diversity in the worker force, we used the black garden ant *L. niger* to experimentally combine workers from one or three source colonies and provided them with an unrelated queen. *L. niger* colonies have a single queen, which in Northwestern Europe is usually mated with a single male, leading to highly relatedness among workers^65^. Queens in this species can also be mated twice or more, but mostly in other geographic regions^18,62^. Established colonies in the field are large, with as much as 10,000 workers^66^, which makes it easy to collect large quantities of brood. After their nuptial flights in summer, hundreds of young mated queens can easily be collected as they roam on the ground looking for a nest site^66^.

### Collection and housing

We collected 44 *L. niger* queens after their nuptial flight on July 10^th^ 2019 on the campus of Johannes Gutenberg University of Mainz, Germany. One day after collection, we transferred each queen to a glass tube half filled with water blocked by cotton and closed with another piece of cotton. Then, we kept the queens in darkness at 21°C and approximately 80% humidity and without food, as *L. niger* founding queens do not feed^67^. These queens had produced a first cohort of at least five workers by the time the experiments began. These workers will hereafter referred to as chaperones.

We collected workers and brood from nine different *L. niger* colonies in the area around the Opel Arena stadium in Mainz, Germany between October and December 2019. In the laboratory, we relocated workers and brood from the soil into glass tubes with water blocked by cotton and covered with aluminum foil. Workers and brood from the same colony were stored in closed boxes (31×22×5cm) coated with fluon in a climate cabinet at 28°C and approximately 100% humidity, and fed five times a week with frozen crickets and a mixture of honey, eggs and vitamins^68^. At the time of collection, these colonies (hereafter referred to as “source colonies”) contained 682 ± 414 (mean ± sd) larvae. We regularly checked the source colonies for pupae to be used for the setup of experimental colonies.

### Setup of experimental colonies

To manipulate worker diversity, we grouped workers produced by either one or multiple source colonies. Because *L. niger* workers are aggressive towards workers from other colonies, we produced experimental colonies by combining pupae, rather than workers, from one or multiple source colonies. The workers that later emerged from those pupae did not aggress each other because they did not differ in their cuticular hydrocarbon profiles, which ants use for nestmate recognition ^69,70,71–73^.

The low diversity experimental colonies were produced by combining 30 pupae from a single source colony and are thus referred to as “control” colonies. We produced the high diversity experimental colonies (hereafter referred to as “treatment” colonies) by combining 30 pupae from three different source colonies (10 pupae per colony). For each experimental colony, we combined the 30 pupae with one unrelated, founding queen and five chaperones to care for the pupae. For each experimental colony, we removed those chaperones, as well as all the eggs present at the time, once three workers had emerged from the pupae. This day was considered as day 0 in the analysis.

We kept the experimental colonies in closed plastic boxes (11×15×3cm) coated with fluon, which contained a glass tube filled with water and cotton as a nest and water source, and a small petri dish for food. We fed the experimental colonies twice a week with frozen crickets and a mixture of honey, eggs and vitamins^68^. From day 0 to day 2, we kept the experimental colonies in a dark climate cabinet at approximately 28°C and 100% humidity. On day 3, we moved the experimental colonies to a climate chamber at 21°C and approximately 80% humidity and in dark conditions.

In total, we set up 43 colonies. We excluded two control and two treatment colonies due to high mortality of pupae and workers. This resulted in 21 control and 18 treatment colonies in the analysis. The experimental colonies varied in how many workers emerged from the 30 pupae that were provided (20.1 ± 5.4, mean ± sd), but this number did not differ between control and treatment colonies (ANOVA: χ^2^ = 1.83, df = 1, p = 0.18).

### Brood production monitoring

In each experimental colony, we counted the number of eggs, larvae and pupae five times a week for 70 days after colony setup. Because the experimental colonies varied in the time to reach day 0 (9 ± 6.6, median ± sd), we only kept the time points between day 0 and day 60 in the analysis to ensure that at any given time point, more than half the experimental colonies were monitored. By the end of the monitoring, only 14 out of 41 colonies had pupae, and even those had a low number of pupae (1.9 ± 2, mean ± sd), and no workers had emerged from the eggs produced in the experimental colonies. Thus we restricted our analysis of brood production to the production of eggs and larvae.

### Foraging assays

We performed the foraging assays 28 days after the last pupa was observed in the experimental colonies to limit age differences across colonies. One treatment colony was not tested because it still contained two pupae at the end of the experiment. Five days prior to the foraging assays, we removed the food to increase the motivation of workers to forage. We performed the foraging assays inside the box of the experimental colonies by placing a small petri dish with a small cotton roll soaked with a honey solution (0.5 ml honey in 1 ml water). Then we observed the colonies for two hours to score the maximal number of workers observed at the food source and to record the time when the first worker arrived at the food.

### Body size measurements

At the end of the experiments, all workers from the experimental colonies were frozen at −18°C for later morphological measurements. To estimate body size, we measured the width of the worker heads as the distance between the outer points of the eyes^64,74–76^. The frozen workers were placed flatly on modeling clay, photographed with a Leica S9i microscope, and measured with LAS V4.12 Leica computer software. We measured all workers that survived the experiment (326 workers) in eight control and eight treatment colonies, while making sure that we only used one experimental colony per source colony.

### Statistical analysis

To test whether control and treatment colonies differed in brood production over time, we built non-linear mixed effect models with the R package *nlme*^77^.

To model the egg production, we used the quintic function

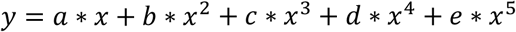

where *y* is the number of eggs, *x* is the number of days, and the parameters *a*, *b*, *c, d* and *e* are estimated by the model to provide the best fit to the empirical data.

To model the larva production, we used a logistic growth equation with the SSlogis() function

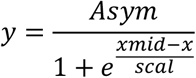

where, *y* is the number of larvae, *x* is the number of days, and the parameters *Asym, xmid* and *scal* are estimated by the model to provide the best fit to the empirical data. *Asym* represents the horizontal asymptote of the logistic growth function, *xmid* the x value of the sigmoid’s midpoint, and *scal* the rate of the logistic growth. The starting values for the parameters were obtained by fitting non-linear models without random effects using the nls() function. For both eggs and larvae, the non-linear mixed effects models were fitted using the function nlme(), and included the worker diversity (control or treatment) as fixed effect and the source colony as random effect. The summary() function was used to extract the estimate for each parameter and each treatment, as well as to test whether estimates differed between treatments. In addition to the non-linear modeling of the change in brood number over time, we used the simpler approach of extracting the maximum number of eggs and larvae recorded in each colony during the experiment. We then tested the effect of worker diversity on this number using the lmer() function with the package *lme4*^78^ to fit a linear mixed effects model with worker diversity as fixed effect and colony as random effect.

We tested the effect of worker diversity on the maximum number of foragers at the food source, as well as the square root transformed data of the time for the first worker to reach the food by building a linear mixed effect model using the lmer() function, with worker diversity as fixed effect and colony as random effect.

To test whether control and treatment colonies differ in size, we calculated the square root of the average head width per colony. To investigate the effect of worker diversity on size variation, we extracted the standard deviation of head size in each colony. Then we built linear models with the lm() function to explain both measurements by worker diversity.

We checked all linear models for normal distribution of the residuals and used the Anova() function of the package *car* ^79^ to test the effect of the explanatory variables. We produced all plots with the packages *ggplot2*^80^ and *ggpubr*^82^, and used the package *dplyr*^81^ for data handling. We ran all analyses in R^83^ version 3.6.1.

## Results

### Experimental increase in worker diversity enhanced the production of larvae, but not eggs

To measure a potential effect of worker diversity on offspring production we monitored the number of eggs and larvae in experimental colonies with low (control) and high (treatment) worker diversity.

The change over time in the number of eggs recorded in the experimental colonies did not differ between control and treatment colonies (p ≥ 0.1 for all parameters; Table 1; Figure 1). Consistently, we could not detect any effect of treatment on the maximum number of eggs recorded in the colonies (ANOVA: χ^2^ = 1.03, df = 1, p = 0.31; Supplementary Figure 1 A).

**Table 1.**
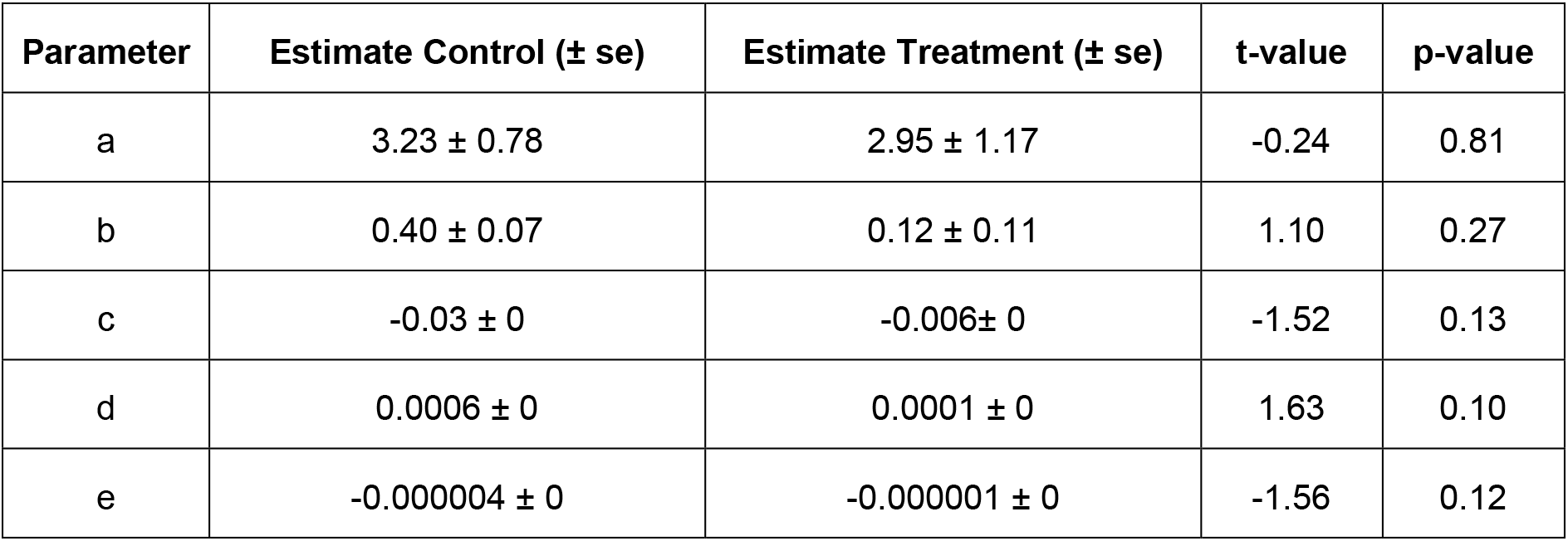
Parameters of the models for egg production over time in the control and treatment colonies. The models are based on a quintic function y = a * x + b * x^2^ + c * x^3^ + d * x^4^ + e * x^5^. y stands for the number of eggs, x is the number of days after setup, and a, b, c, d and e are the parameters estimated by the models.

| Parameter | Estimate Control (± se) | Estimate Treatment (± se) | t-value | p-value |
| --- | --- | --- | --- | --- |
| a | 3.23 ± 0.78 | 2.95 ± 1.17 | -0.24 | 0.81 |
| b | 0.40 ± 0.07 | 0.12 ± 0.11 | 1.10 | 0.27 |
| c | -0.03 ± 0 | -0.006 ± 0 | -1.52 | 0.13 |
| d | 0.0006 ± 0 | 0.0001 ± 0 | 1.63 | 0.10 |
| e | -0.000004 ± 0 | -0.000001 ± 0 | -1.56 | 0.12 |

**Figure 1.**
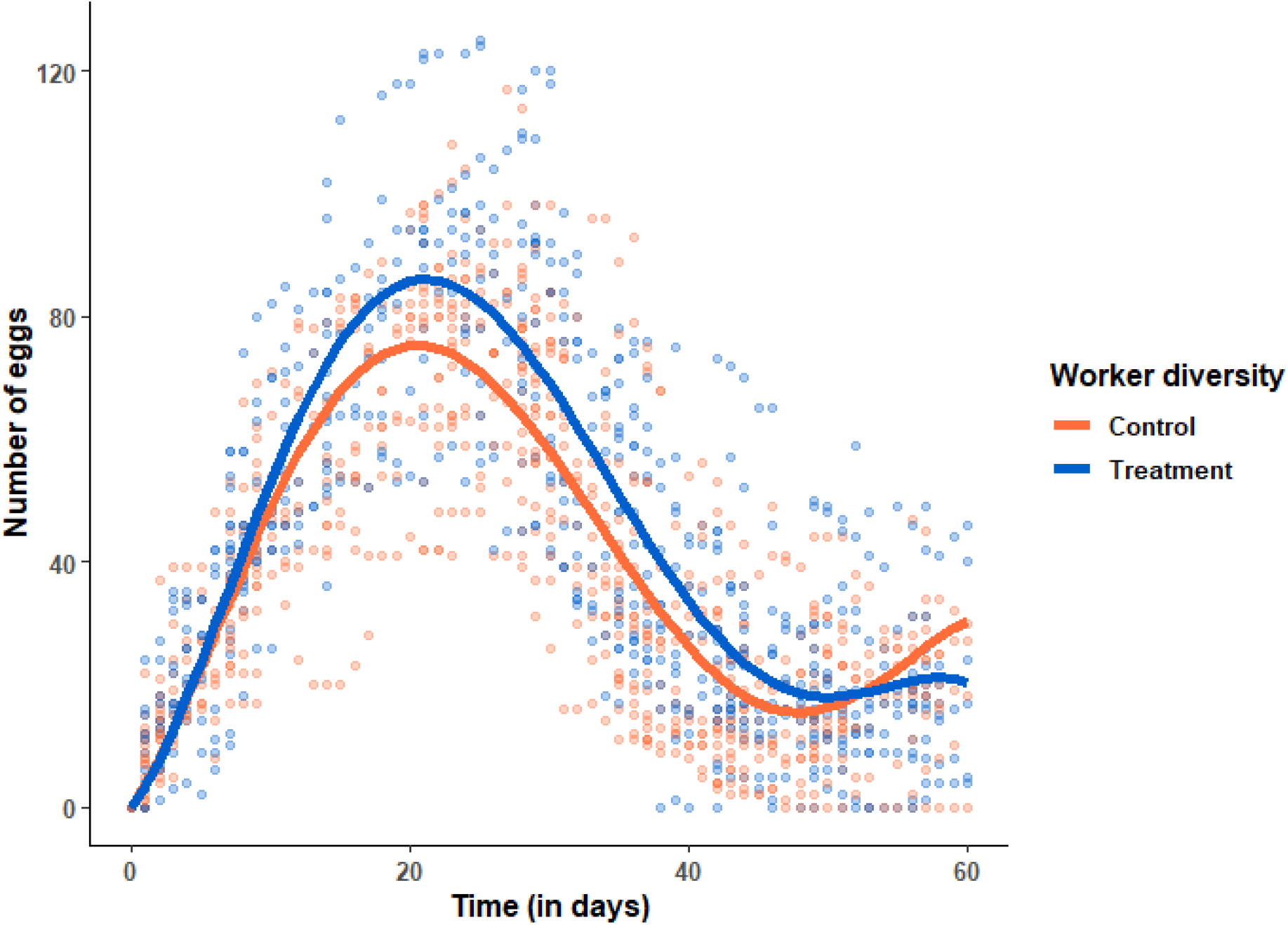
Number of eggs in control and treatment colonies over time. The dots show the raw data for all colonies and time points. The curves depict the output of the models for control (orange) and treatment (blue) colonies. None of the parameters of the quintic function differed significantly between control and treatment colonies (all p ≥ 0.1; Table 1).

The change over time in the number of larvae recorded in the experimental colonies differed significantly between control and treatment colonies (Table 2; Figure 2). Specifically, we found that colonies with higher worker diversity reached a higher horizontal asymptote by the end of the experiment (*Asym*, t = 2.56, p = 0.011; Table 2; Figure 2), and showed a higher logistic growth rate (*scal*, t = 2, p = 0.046; Table 2; Figure 2). We did not detect any effect of worker diversity on the timing of the logistic growth (*xmid*, t = 0.94, p = 0.35; Table 2; Figure 2).

**Figure 2.**
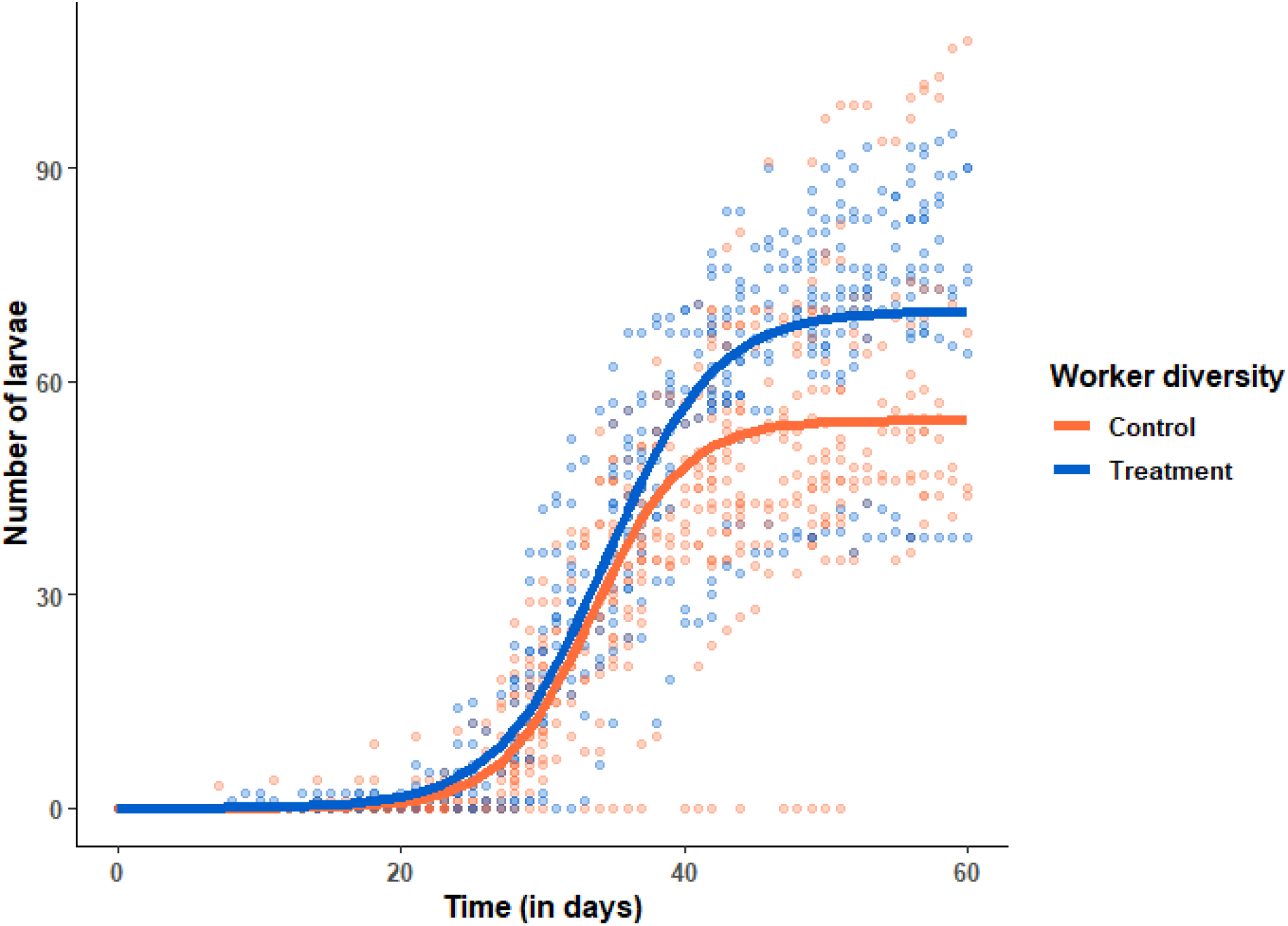
Number of larvae in control and treatment colonies over time. The dots show the raw data for all colonies and time points. The curves depict the output of the models for control (orange) and treatment (blue) colonies. Two parameters of the logistic growth function differed significantly between control and treatment colonies (Table 2).

**Table 2.** Parameters of the models for larva production over time in the control and treatment colonies. The models are based on a logistic growth function 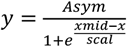. y stands for the number of larvae, x is the number of days seal after setup, and the parameters estimated by the models are the asymptote (Asym), the timing of the growth (xmid) and the rate of the growth (scal).

| Parameter | Estimate Control (± se) | Estimate Treatment (± se) | t-value | p-value |
| --- | --- | --- | --- | --- |
| *Asym* | 54.6 ± 4.0 | 70.0 ± 6.0 | 2.56 | 0.011 |
| *xmid* | 33.5 ± 0.7 | 34.5 ± 1.0 | 0.94 | 0.349 |
| *scal* | 3.24 ± 0.2 | 3.9 ± 0.3 | 2.00 | 0.046 |

In one control colony, no larvae were produced throughout the experiment (Figure 2). To ensure that the effect of worker diversity on larva production did not stem from this colony only, we repeated the analysis after excluding this colony and found qualitatively similar results (*Asym*, t = 2.36, p = 0.018, *xmid*, t = 0.77, p = 0.44, *scal*, t = 1.87, p = 0.061). We also found that the maximum number of larvae recorded in each experimental colony was higher in treatment than in control colonies (ANOVA: χ^2^ = 4.87, df = 1, p = 0.027, Supplementary Figure 1 B).

### Experimental increase in worker diversity did not affect foraging

We could not show that control and treatment colonies differed in the maximum number of workers at the food (ANOVA: χ^2^ = 2.05, df = 1, p = 0.15, Figure 3 A), nor in the time for the first worker to reach the food (ANOVA: χ^2^ = 0.11, df = 1, p = 0.74, Figure 3 B).

**Figure 3.**
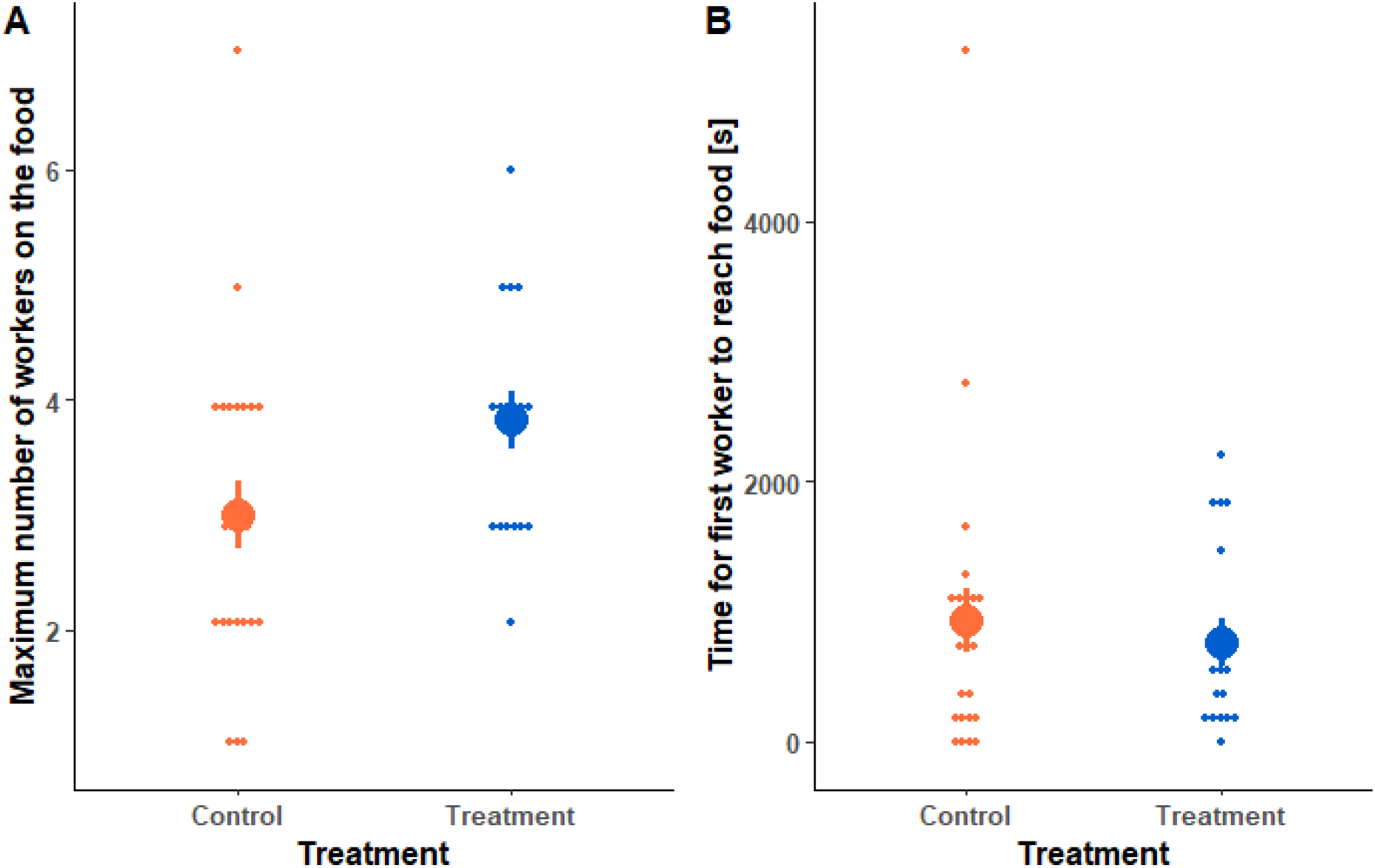
Foraging did not differ between control and treatment colonies. The large dots depict the mean ± standard error. The small dots represent the raw data points. We did not detect any significant difference between control and treatment colonies in (A) the maximum number of workers (ANOVA: χ^2^ = 2.05, df = 1, p = 0.15) and (B) the time for the first worker to reach the food (ANOVA: χ^2^ = 0.11, df = 1, p = 0.74).

### Experimental increase in worker diversity enhanced variation in body size

We extracted the standard deviation in head width for each experimental colony to find that this measure of body size variation differed between control and treatment colonies, as colonies with higher worker diversity showed higher variation (ANOVA: F_1,14_ = 26.42, p < 0.001, Figure 4 A). However, we did not find such an effect of worker diversity on the average mean head width per colony (ANOVA: F_1,14_ = 0.099, p = 0.76, Figure 4 B).

**Figure 4.**
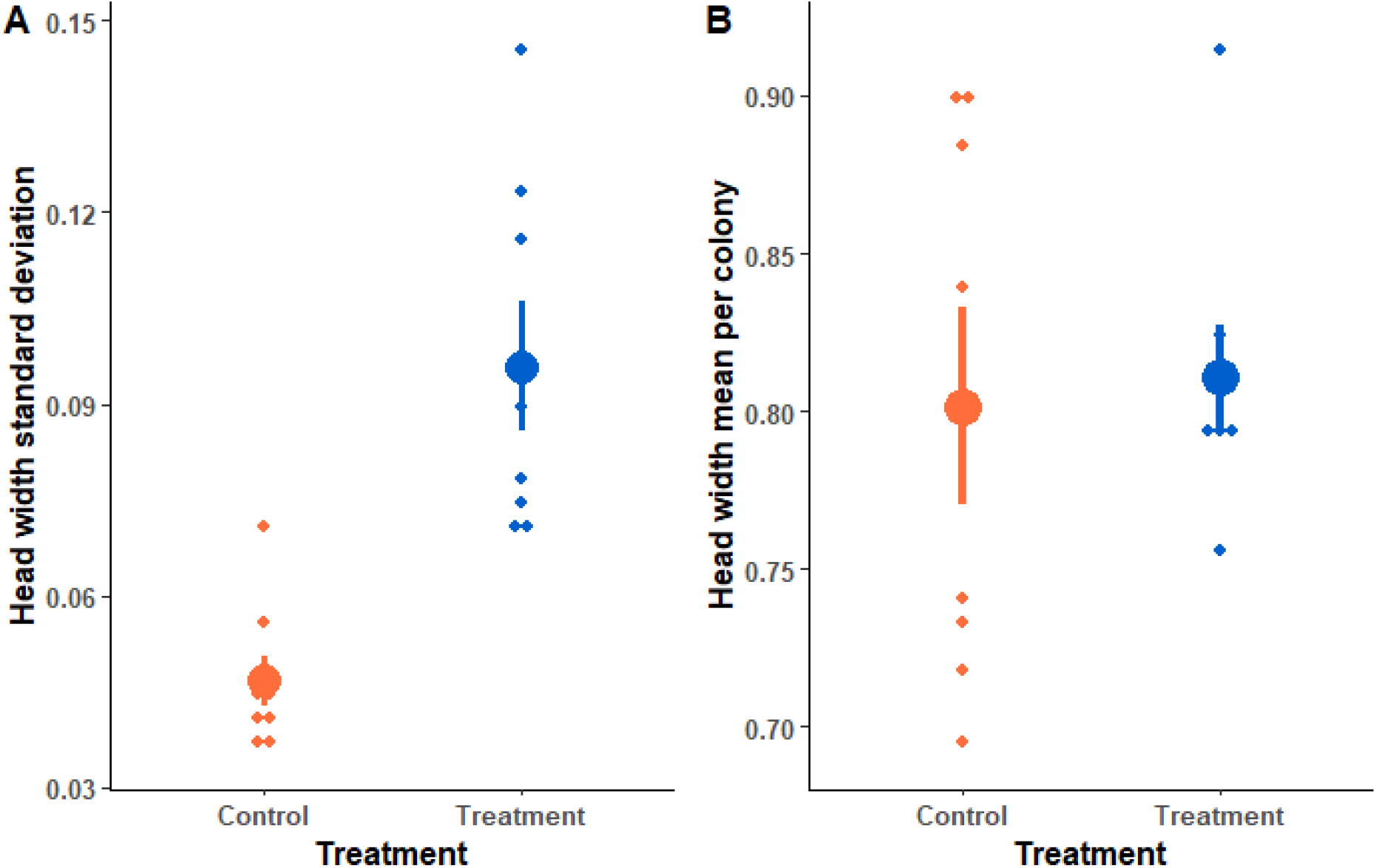
Control and treatment colonies differed in the standard deviation, but not the average, in head width. Variation. The large dots depict the mean ± standard error. The small dots represent the raw data points. The experimental increase in worker diversity resulted in (A) a higher within-colony variation in head width (ANOVA: F_1,14_=26.42, p < 0.001) but (B) no difference in the average head width per colony (ANOVA: F_1,14_ = 0.099, p = 0.76).

## Discussion

The aim of this study was to test the effect of worker diversity on colony performance in eusocial insects. To do so, we experimentally increased worker diversity of *Lasius niger* colonies by combining workers form three different source colonies and compared them to colonies composed of workers from a single source colony. In addition, we provided the experimental colonies with unrelated queens to disentangle any effects of worker diversity from maternal effects. We found that increased worker diversity enhanced the production of larvae, as well as size diversity within the worker force. We did not detect any effect of worker diversity on egg production and foraging performance.

Our finding that worker diversity enhanced larva production in experimentally produced laboratory colonies is consistent with previous reports of benefits provided by higher genetic diversity in other species of eusocial Hymenoptera^19,24,36,48,55–58^. In the honey bee *Apis mellifera*, decreased genetic diversity by artificial insemination, or restricted natural mating was shown to result in lower productivity and fitness^24,48,58^. Similarly, in bumble bees (*Bombus terrrestris*) higher genetic variance was shown to decrease parasite load and enhance reproductive success^20,56,57^. Our study adds to previous reports because we found benefits of experimentally increasing worker diversity in a species with lower natural levels of diversity, while controlling for any maternal effects caused by the experimental manipulation. Assuming that larva production is a relevant proxy for colony performance and fitness, our study validates the hypothesis that worker diversity positively affects colony fitness.

Worker diversity may improve colony performance via a more efficient division of labor^41^. Behavioral division of labor among workers likely stems from workers differing in their response thresholds to perform specific tasks^32^. In our study, the experimental colonies produced with workers from three source colonies were more diverse than the control colonies in terms of genetic background and size variation. This is expected, as genetic effects on worker size and morphology have been reported in multiple species of ants ^84–86^ and worker size differs among colonies in *L. niger*^87^. Worker size and genetic background both influence the response threshold of individual workers^28,84,88–90^, and worker size polymorphism is generally associated with improved division of labor^91–93^. In this perspective, a more diverse worker force would have resulted in a more heterogeneous mix of response thresholds, possibly enhancing the efficiency of division of labor among workers^36–39,94–96^.

The beneficial effect of worker diversity on the production of larvae likely stemmed from more efficient brood care. We detected an increase in the number of larvae, but not in the number of eggs, and we could not detect any effect on foraging. This suggests that increased worker diversity improved the survival and/or development of larvae, but probably not via better food provisioning to the brood. It may be that our experiment was unlikely to detect a difference in foraging because of the low number of foragers, but if more diverse colonies were better at foraging for food, we would have expected the better nourished queens in those colonies to produce more eggs. Our results do not support this expectation. Another explanation that could explain our inability to detect a difference in egg number is egg cannibalism by the larvae, which is common in eusocial Hymenoptera^2,97,98^. More diverse colonies had more larvae, which in turn may have eaten more eggs compared to low diversity colonies. According to this scenario, the number of eggs would differ between more and less diverse colonies before the first larva emerged from the eggs. This was not the case, as the maximum number of eggs – which was reached before larvae appeared – was not affected by worker diversity. Our findings are more consistent with worker diversity improving brood care, although we do not have direct evidence for such an effect.

A minor proportion of previous studies did not detect an association between diversity and colony performance^50,60,61,63,64,74,99,100^ or had ambiguous findings^55^, including in *L. niger* field colonies^46,62^. Discrepancies among studies suggest that the effect of diversity on division of labor and colony fitness is context- and/or species-dependent. In our study, we used an experimental approach to control for confounding factors, but in doing so, we produced an artificial situation of small colonies with same-age workers and an unrelated queen in the laboratory. While this does not resemble a natural situation, our results provide robust evidence that increased worker diversity provides benefits in some situations. We did not manipulate genetic diversity but combined workers from multiple source colonies to produce diversity in the worker force. We confirmed that our experimental manipulation increased size variation in the more diverse colonies. Such size diversity could stem from genetic differences across colonies, but could also be explained by environmental and maternal effects^84,85,87,101^. The genetic background affects size and morphology in eusocial insects^84–86^, thus one strategy to increase worker size diversity is to increase genetic diversity. Our finding that worker size diversity is linked to colony performance in some conditions could explain why otherwise costly strategies to increase genetic diversity (e.g., multiple mating and/or multiple queens per nest) have been selected in some eusocial insect species^5,11,53,102^.

So far, evidence for benefits of worker diversity in eusocial insects came from correlative studies, experimental studies where low diversity was also the unnatural situation and/or where other confounding factors such as maternal effects could have played a role. In this study, we experimentally increased worker diversity and controlled for maternal effects. We found that increased worker diversity improved larva production, a proxy for colony performance and fitness, possibly via enhanced division of labor. Our findings confirm that increased diversity can benefit colony fitness in some situations, which could have led to the evolution in some eusocial insects of multiply mated queens and multiple-queen colonies.

## Supplement

**Supplement Figure 1.**
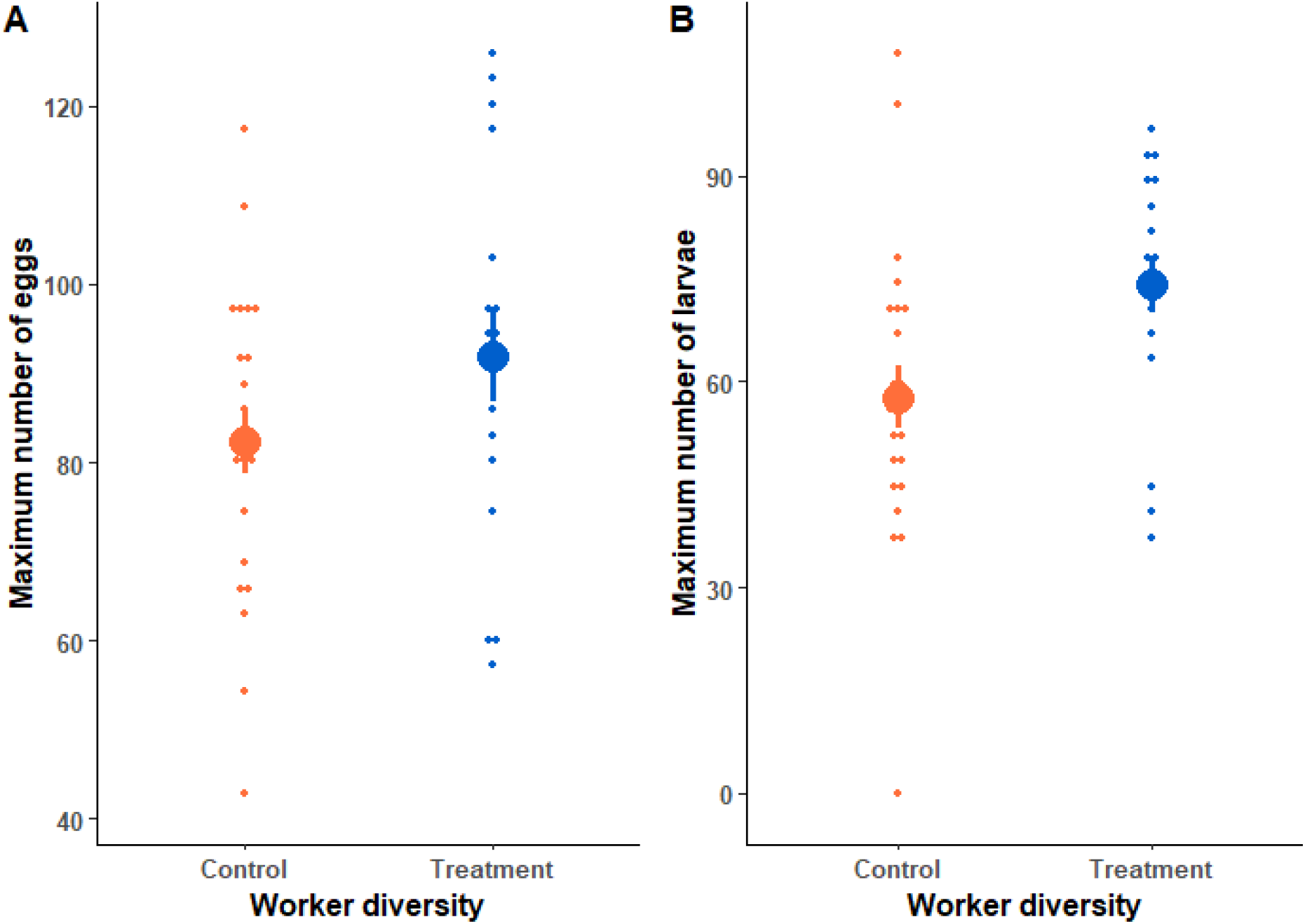
Maximum number of brood items in the different treatments. Maximum number of (A) eggs and (B) larvae produced by control and treatment colonies within 60 days of monitoring. (A) There is no significant difference between the maximum number of eggs produced between treatments (ANOVA: χ^2^ = 1.03, df = 1, p = 0.31). (B) The maximum number of larvae between treatment and control colonies was significantly different (ANOVA: χ^2^ = 4.87, df = 1, p = 0.027).

